# NOVEL CHARACTERISTICS OF SMALL NONCODING RNAS IN RAT EPIDIDYMAL EXTRACELLULAR PARTICLES

**DOI:** 10.1101/2022.05.27.493742

**Authors:** Ross Gillette, Dana L. Sheinhaus, Shyamal Waghwala, Andrea C. Gore

**Author notes:** **Author Contributions:** Conceptualization: R.G.; Methodology: R.G., D.L.S.; Investigation: R.G., D.L.S.; Formal Analysis: R.G., S.W.; Writing – Original Draft: R.G.; Writing – Review & Editing: R.G., D.L.S., A.C.G.; Funding Acquisition: R.G., A.C.G.

## Abstract

Extracellular vesicles (EVs) released from the epididymal epithelium, referred to as epididymosomes, impart functional competence on sperm as they transit the epididymis by merging with sperm and releasing a complex repertoire of molecules. The cargo of epididymal EVs includes small noncoding RNAs (sncRNAs) that are modified by external factors such as stress, nutrition, and drug use, which are delivered to sperm and affect offspring development and outcomes. The rat is an important translational model system for many fields including reproduction and transgenerational epigenetics, but there is little work on rat epididymosomes and their sncRNA cargo. To fill this gap in knowledge, in the current study we performed a comprehensive characterization of an epididymal EV preparation that we refer to as extracellular particles (EPs) because of the potential presence of other non-vesicular extracellular particles along with the EVs. EPs were collected from the caput epididymis, isolated, subjected to verification of the presence of EVs, and sncRNAs were sequenced. We considered nearly all known categories of small RNA and their fragment sub-types and in the process identified a subset that did not strictly fit the definition of any known category. These unique small RNAs are expressed strictly from within the boundaries of CpG islands and have a distinct 19-nt length. Their functional significance remains unknown, but they have characteristics of RNA fragments that can associate with the Argonaute/PIWI family of proteins and therefore could have regulatory function via RNA induced silencing or de novo DNA methylation.

## Introduction

Spermatozoa are incapable of fertilization as they exit the testes [1] and only gain motility and the capacity for fertilization as they transit the epididymis from the caput to the cauda [2]. This process requires epididymosomes [3–6], a type of extracellular vesicle (EV) within the microvesicle category [7–9]. Epididymosomes are released from the epididymal epithelium via apocrine secretion [10], fuse to spermatozoa [11], aid in their maturation and functionalization [4,12,13], and mark dead or dying spermatozoa for elimination [14,15]. Epididymosomes carry a complex repertoire of ions, proteins [16], phospholipids [17], metabolites [18], and small non-coding RNAs (sncRNA) [4,12,19,20] that are transferred to [21,22] and alter the function of the spermatozoa or impart functional consequences on the fertilized embryo [23]. Importantly, sncRNA molecules transferred to spermatozoa have the capacity to alter the epigenetic landscape of either the spermatozoa or the fertilized embryo and thereby influence subsequent generations [22,24]. Furthermore, the sncRNA contents of epididymosomes are subject to modification by the hormonal milieu [25,26] and environmental challenges such as psychosocial or metabolic stress [26–31], making them a potential vector for intergenerational effects in offspring. This key role was illuminated by studies showing that early life stress caused behavioral and physiological changes in the offspring, phenotypes that were recapitulated by microinjection of a small number of microRNAs into a fertilized egg subsequently used for *in vitro* fertilization [24,32,33].

The sncRNA repertoire of epididymosomes is complex and diverse, with most information in model systems from mice [34]. The most abundant sncRNA found in mouse epididymosomes is microRNA (miRNA) followed by smaller proportions of transfer RNA (tRNA) and ribosomal RNA (rRNA) [29,34,35]. The remainder consists of piwi-interacting RNA (piRNA), small nuclear RNA (snRNA), and small nucleolar RNA (snoRNA) [29]. Other species of sncRNA, such as Y RNA and Vault RNA, have not yet been reported in epididymosomes but should not be ruled out for inclusion in these complex profiles. Furthermore, there are likely additional sncRNAs yet to be identified [36].

Much of the knowledge we have on the sncRNA contents of caput epididymosomes, which are crucial for the final steps of spermatozoa maturation, is from mouse models [29,34,35]. However, there is minimal conservation between the miRNA in caput EVs between mice and humans [37] and to our knowledge, other classes of sncRNAs have not been directly compared between species. Very little has been reported on the epididymosomes of rats, which are an important model system for endocrinology, reproduction, and epigenetic transgenerational inheritance, and in which behavioral work is considered to better translate to humans [38–40]. Here, we characterized the contents of EPs derived from the caput epididymis of the male rat, during which we identified what we believe to be a previously undefined class of sncRNA.

## Materials and Methods

### Animals and treatment

All animal experiments were conducted using humane procedures that were approved by the Institutional Animal Care and Use Committee at The University of Texas at Austin and in Accordance with National Institutes of Health (NIH) and Animal Research: Reporting of *In Vivo* Experiments (ARRIVE) guidelines [41]. Three-month old male and female Sprague Dawley rats were purchased (Envigo, Indianapolis, IN) and shipped to the Animal Resource Center at the University of Texas at Austin and allowed two weeks to acclimate to the housing facility. All animals in the colony were housed in a room with consistent temperature (∼22° C) and light cycle (14:10 light:dark), had *ad libitum* access to filtered tap water and a rat chow with minimal phytoestrogens (Teklad 2019: Envigo, Indianapolis, IN), and housed in cages lined with wood-chip bedding (Sani-Chips: P.J. Murphy, Montville, NJ). After acclimation, female rats were observed for vaginal cytology as previously described [42,43] indicating proestrus, and therefore receptivity. Receptive female rats were randomly paired with an experienced breeder male, receptivity was confirmed via the observation of lordosis during a mount, and the pair was left together overnight. The following morning, vaginal cytology was checked for the presence of sperm and if present, the animal was marked as embryonic day (E1). The pregnant dams were single-housed and provided nesting material on E18.

On E8, the dams (N = 6) were randomly split into two groups (each N = 3) and fed ¼ Nilla Wafers™ daily with either 3% DMSO or 1 mg/kg Aroclor 1221 (A1221) (#C-221N, Accustandard, Lot #072-202-01 - an estrogenic polychlorinated biphenol [44]) from E8 – E18 and postnatal day (PND) 1 – 21. The original goal of work was to characterize sncRNAs in rat epididymal EVs, as well as to generate pilot data on effects of the endocrine-disrupting chemical, A1221. However, no differential expression of sncRNAs was caused by treatment after corrections for multiple comparisons. Therefore, treatment was not considered or analyzed in any of the data presented in this manuscript with current work focused solely on characterization of the composition of EV sncRNAs. Litters were culled to 5 females and 3 males based on median anogenital index at P1. At PND 21, all pups were weaned and housed in cages of two or three same-sex animals. Only males were used in the present manuscript and were otherwise unmanipulated until euthanasia besides weekly handling to acclimate each rat to the experimenters in order to reduce stress. Females were allocated to other projects.

### Sample Collection

At PND 105, one male rat was randomly selected from each of the 6 litters using a random number generator (N = 1 per litter) and euthanized by rapid decapitation. The testis and epididymis were removed via a small incision in the scrotum. The epididymis was separated from the testis and segmented into three portions (caput, corpus, and cauda). The caput of the epididymis was minced into small pieces with scissors and placed in warm (37° C) M2 culture media (M7167, Millipore Sigma) with HEPES (5.43 g/L) and 2% exosome depleted fetal bovine serum (A2720803, ThermoFisher). The slurry was placed on a rocker at room temperature for 30 minutes to allow sperm and epididymal fluid to suspend in solution, the supernatant was removed and large tissue chunks were excluded. The resulting supernatant was centrifuged at 500 X g for 5 minutes to pellet and remove sperm. The supernatant was again removed, immediately frozen on dry ice, and stored at -80° C until use (∼ 3 mo) which is the preferred method of storage when necessary [45] and within the timeframe before which substantial effects to EVs are observed [46].

### Extracellular Particle Purification and Nucleic Acid Extraction

The buffer supernatant containing extracellular particles from the caput of the epididymis was thawed on ice for 30 minutes prior to isolation. When fully thawed the buffer was mixed by pipetting and then filtered with a 0.8 μm syringe filter (Millex-AA, SLAA033DD, Millipore Sigma) to remove any cells or large cellular debris. Extracellular particles and RNA were sequentially isolated from the media using the Qiagen exoRNeasy Midi kit [47] (#77144, Qiagen, Germantown, MD) according to the manufacturer’s protocol. Briefly, 200 μl of media from each sample was passed through a spin column that selectively binds extracellular vesicles. The column was washed to remove contaminants and debris, and then eluted in Qiazol lysis buffer (#79306, Qiagen, Germantown, MD). Chloroform (AAJ67241AP, FischerScientific) was added, mixed by shaking, and incubated at room temperature to allow phase separation. Phase separation was aided by centrifugation at 12,000 X g for 15 minutes at 4° C and the aqueous phase was aspirated and passed through a second membrane spin column that selectively binds RNA. The membrane was washed to remove contaminants and the sample eluted in 13 ul RNase free water. M2 culture media and 2% exosome depleted fetal bovine serum was used as a negative control to determine exogenous contamination and subjected to the same extracellular vesicle and RNA isolation procedures described above. These negative controls were analyzed via particle analysis and for resulting nucleic acids to ensure there were no exogenous or contaminating exosomes or nucleic acids. None were found (**Supplemental Figure S1A**).

### Extracellular Particle and RNA Quality Control

An aliquot of the extracellular particles from the same samples described above were isolated using the exoEasy Maxi Kit (76064, Qiagen) according to the manufacturer’s protocols. The resulting EPs from two samples were analyzed using a NanoSight 300 (Malvern Panalytical) in 5 replicates to establish a size distribution of the resulting EPs and to ensure there were no contaminating particles. RNA extracted from isolated EPs was first quantified (ND-1000, ThermoScientific) and diluted for size distribution analysis and quality control using the small RNA (5067-1548, Agilent) and pico RNA kits (5067-1514, Agilent) on a BioAnalyzer 2100 (Agilent). We found there was no cellular contamination indicated by a lack of 18s and 28s ribosomal RNA [48,49] and that the majority (∼80%) of extracted RNA was in the 20-40 bp range, indicative of small RNA molecules (**Supplemental Figure S1B**).

### Characterization of Extracellular Particles by Electron Microscopy

Transmission electron microscopy (TEM) and cryo-electron microscopy (cryo-EM) were performed at the University of Texas’s Center for Biomedical Research Support (CBRS). TEM specimens were pre-fixed in microfuge tubes by adding 0.1 μl of electron microscopy grade 16% paraformaldehyde (Electron Microscopy Sciences) to 5 μl purified specimen and incubating for 20 minutes at room temperature. Negative staining was performed by applying 5 μl of pre-fixed sample to a glow discharged grid and allowing to adhere for 1-2 minutes, then washed on a drop of molecular grade water, stained with 2% aqueous uranyl acetate, blotted, and allowed to dry prior to imaging in an FEI BioTwin electron microscope fitted with an AMT 1kx x 1kx camera.

Cryo-EM specimens were applied to a glow-discharged lacey carbon grid. The grid was incubated for 30 seconds at 4° C and blotted with filter paper to leave only a thin film spanning the grid holes. The sample was kept at 100% humidity before plunge-freezing into liquid ethane using a Vitrobot (CBRS, UT Austin). The vitreous sample grids were observed on a Glacios microscope (Thermo Fisher) and images were captured at 200 kV with a 1.97 Å^2^/pixel Falcon 4 camera. Data were collected using SerialEM version 3.9.0 beta and motion corrected using cryoSPARC with 60 frames.

### Library Preparation, Sequencing, and Quality Control

Library preparation was performed at the University of Texas Genomic Sequencing Facility (GSAF) using the NEBNext Small RNA library preparation kit (E7330, New England Biolabs) as previously published [50]. Samples were prepared with 14 cycles of PCR and the final product was size selected using a 3% gel cassette on the Blue Pippin instrument (Sage Sciences), with the parameters set to 105-165 bp. Final size selected libraries were checked for quality on the Agilent BioAnalyzer with High Sensitivity DNA analysis kit (5067-4626, Agilent) to confirm proper size selection. The Kapa Library Quantification kit for Illumina libraries (KK4602) was used to determine the loading concentrations prior to sequencing. Samples were sequenced on a NovaSeq 6000 Single Read, SR50 run, with read counts ranging from 25 to 32 million per sample. The raw sequence data is publicly accessible (GSA: CRA008039).

### Analysis Pipeline

Raw RNA reads were first passed through quality control (FastQC) and checked for read quality and size distribution, which showed excellent average read quality (avg. PHRED > 36) and expected read length (50 bp). The raw reads were trimmed for Illumina adaptors (QuasR, BioConductor) and again passed through quality control (FastQC) to determine per base read quality and size distribution, which again showed excellent average read quality (avg. PHRED > 36) and an expected read length distribution (∼30 nt) based on electrophoresis performed before library prep. The trimmed reads were aligned (RBowtie, Bioconductor) to the unmasked rat genome (Rnor v6) with strict matching parameters to ensure reads were aligned to best-hit locations but were allowed multiple mapping locations (n = 12) due to the promiscuous nature of small RNA origins. If the alignment parameters were met but a read mapped to multiple locations, the read was randomly assigned to one of the similar best hit locations. Alignment efficiency was analyzed for cross-species contamination or PCR primer amplification artifacts and was found to be highly efficient (∼85%). The aligned reads were assigned a feature annotation (Seqmonk, R, BioConductor), visualized, and analyzed for read count and length distribution. Our annotation pipeline considered reads aligned to miRNA, piRNA, rRNA, snoRNA, snRNA, tRNA, long non-coding RNA (lncRNA), vault RNA, and Y RNA. CpG islands were inadvertently included in the pipeline, although at the time we did not have any expectation of finding sncRNAs aligned with this category. We also obtained annotation tracks for nested and simple repeats (UCSC genome database) and microsatellites (UCSC genome database). Reads were assigned an annotation designation only after considering all other annotation types co-occurring in the same region.

## Results

### The size distribution of rat caput epididymosomes

Particle tracking (NanoSight 300, Malvern Panalytical, Malvern, UK) was used to determine the size distribution of the extracellular particles isolated from the caput of the rat epididymis using the Qiagen exoEasy Maxi kit as described above. The isolated EPs from two separate samples were analyzed 5 times each. The first sample had a mean EP size of 181.4 nm (standard error [SE] 3), mode of 152 nm (SE 2.5), and standard deviation of 40.6 nm (SE 1.1) and the second a mean of 178.7 (SE 0.5), mode of 156.1 (SE 3.8), and standard deviation of 45.7 nm (SE 1.8). The size distribution and density of the extracellular particles analyzed are shown in **Figure 1**. Approximately ∼93% of all particles tracked were between 50 and 250 nm in size, which are characteristic of epididymosomes [4].

**Figure 1.**
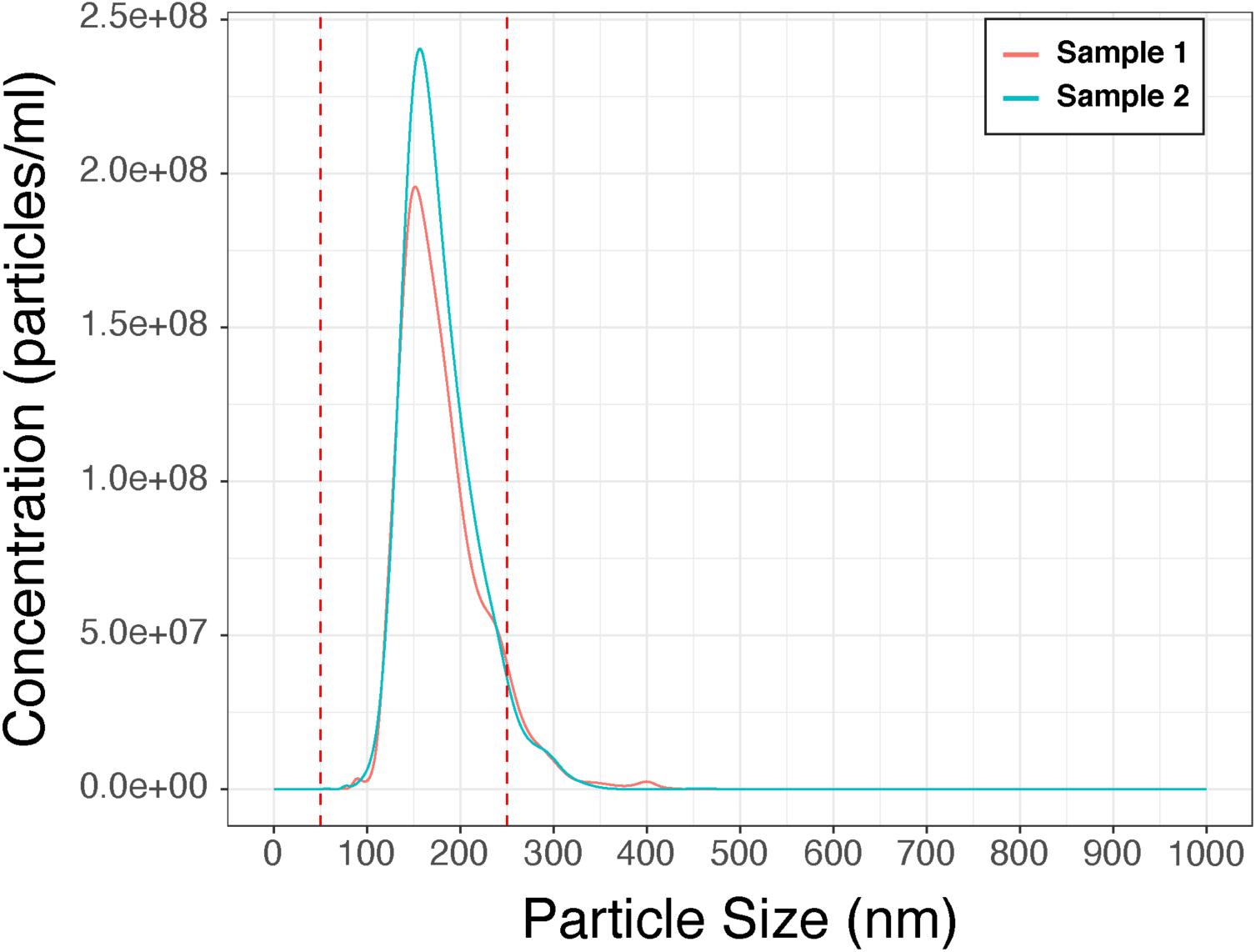
The size distribution and density of the extracellular vesicles from two separate samples are shown as analyzed by nano-particle tracking (NanoSight 300). The red dashed lines indicate the boundaries of size for canonical epididymosomes (50 – 250 nm [4]).

### The morphology of rat caput EPs

TEM and cryo-EM were used to determine the morphology of the extracellular particles isolated from the caput of the rat epididymis using the Qiagen exoEasy Maxi kit as described above. To increase the concentration of vesicles present in the sample for imaging, the epididymis was thoroughly minced with scissors, and caput biofluid was collected by squeezing the tissue with forceps. The isolated EPs from two separate samples were imaged using either TEM or cryo-EM. TEM allowed us to visualize the composition of the sample, whereas cryo-EM aided in visualization of EVs, including lipid bilayers and internal structures. Both types of samples were composed of a population of intact round particles that possess a clear lipid bilayer/membrane, which is characteristic of extracellular vesicles. Single, double, and multilayer vesicles, characteristic of EV preparations and defined by the presence of one, two, or multiple membranes, respectively, are presented in **Figure 2**. Cryo-EM imaging revealed clear lipid bilayer membranes (**Figure 2C, D)**.

**Figure 2.**
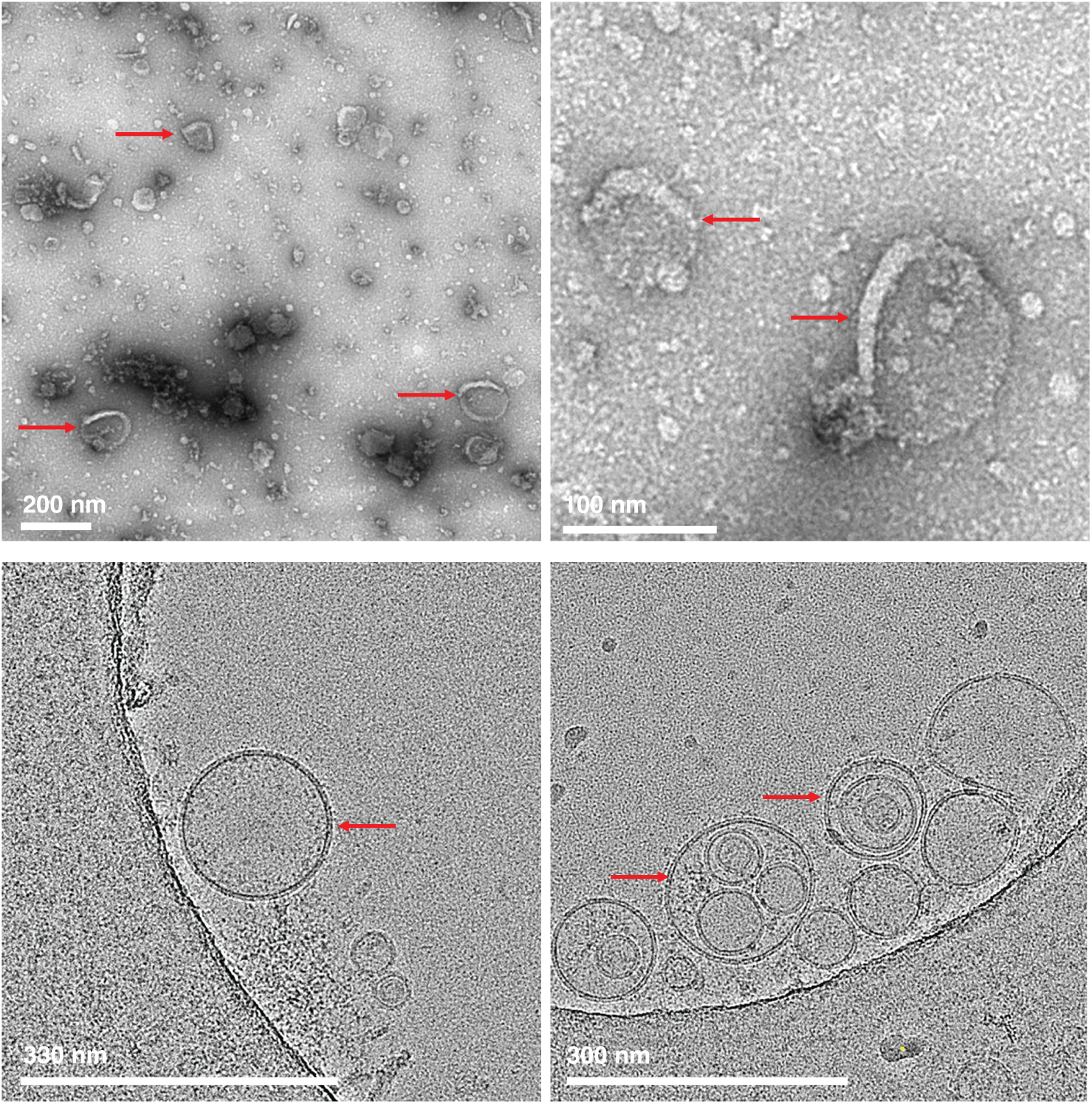
Results from imaging EPs from the rat caput are shown. **A, B)** Representative TEM images of our EP preparation. Red arrows point to several cup-shaped vesicles, typical of EV morphology. **C)** Cryo-EM image of our EP preparation with an arrow pointing towards a single vesicle. **D)** Arrows point towards multilayer vesicles.

### The small RNA contents of rat caput EPs

The sncRNA contents of rat epididymosomes were dominated by reads that aligned to tRNA (79.1%) or piRNA loci (18.1% - **Figure 3A; Tables 1 & 2**). We were surprised at the relatively low abundance of miRNA in rats considering that mice caput EVs primarily consist of miRNA (∼60% -- [20,29,34,35]). All other annotation types, including miRNAs, accounted for the remaining 2.8% of the aligned reads. When tRNA and piRNA reads were removed from analysis and aligned reads were analyzed as a proportion of the remaining reads (**Figure 3B**), the largest remaining category belonged to reads that aligned to CpG islands (55.2%), which is not a known unit of sncRNA expression (discussed below). The remaining reads aligned to Y RNA (24.6%), lncRNA (7.5%), miRNA (6.4%), rRNA (4.3%), and snoRNA (1.8%).

**Figure 3.**
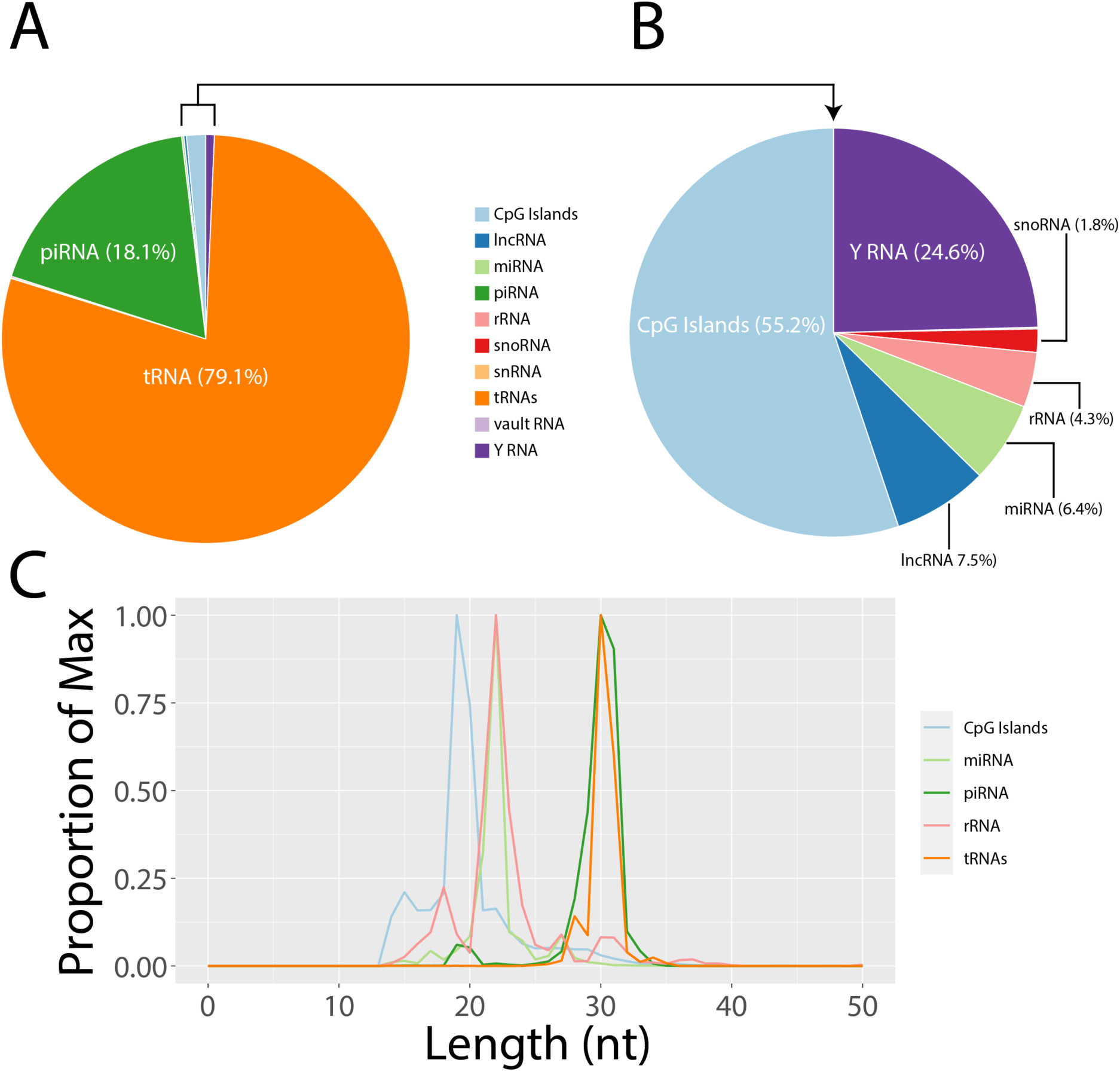
Results from sequencing small RNAs derived from EVs in the rat caput are shown. **A)** The proportion of small RNA reads assigned to each of 10 annotations in our analysis; tRNA and piRNA account for the 97.2% of small RNA in caput EVs while miRNAs only account for 0.177%. **B)** The remaining small RNA reads are shown as a proportion of the residual 2.8% not assigned to tRNA or piRNA loci. CpG islands, which are not a known unit of small RNA expression, account for the majority (55.2%) of the remaining reads. **C)** The size distribution of 4 well-defined sncRNAs are shown together with the one undefined category (CpG islands); the latter has a unique size profile (19 nt). The size distribution for piRNA (29-31 nt) and miRNA (22 nt) are exactly as expected [51,52]. Reads aligned to tRNA (30 - 31 nt) and rRNA (primary peak at 22 nt and small peaks at 18 nt and 30 nt) both demonstrate expected size specific fragmentation [65,103].

**Table 1.** Descriptive statistics of the proportion and read length for each annotated small RNA category are shown, with the mode, median, mean, standard deviation, and length range from which 90 percent of the reads are found. Data are a composite of the 6 individuals.

| Class | % of Total | Mode | Median | Mean | SD | 90% Range (nt) |
| --- | --- | --- | --- | --- | --- | --- |
| tRNAs | 79.124 | 30 | 30 | 30.19 | 1.18 | 28 - 31 |
| piRNA | 18.125 | 30 | 30 | 29.6 | 2.5 | 26 - 32 |
| CpG Islands | 1.518 | 19 | 19 | 19.89 | 3.75 | 15 - 28 |
| Y RNA | 0.676 | 31 | 31 | 28.33 | 3.89 | 22 - 31 |
| lncRNA | 0.207 | 22 | 22 | 22.27 | 4.98 | 15 - 33 |
| miRNA | 0.177 | 22 | 22 | 22.04 | 2.34 | 19 - 27 |
| rRNA | 0.12 | 22 | 22 | 22.65 | 4.06 | 17 - 31 |
| Vault RNA | 0.001 | 18 | 19 | 22.99 | 8.02 | 14 - 41 |
| snRNA | 0.003 | 16 | 16 | 18.05 | 3.85 | 15 - 26 |
| snoRNA | 0.051 | 27 | 24 | 23.44 | 5.12 | 15 - 32 |

**Table 2.** Distribution of the percentages represented by the small RNA categories in the six individual rat caput epididymis EP samples (individual rat identifiers, with the prefix OPA, are provided). There was considerable consistency across the six individuals.

| **Class** | **All**<br>**(%)** | **OPA16A**<br>**(%)** | **OPA21C**<br>**(%)** | **OPA24A**<br>**(%)** | **OPA31A**<br>**(%)** | **OPA22A**<br>**(%)** | **OPA23B**<br>**(%)** |
| --- | --- | --- | --- | --- | --- | --- | --- |
| **tRNAs** | 79.124 | 76.084 | 78.923 | 80.168 | 81.334 | 79.366 | 78.128 |
| **piRNA** | 18.125 | 20.219 | 18.729 | 17.272 | 16.739 | 17.463 | 18.705 |
| **CpG islands** | 1.518 | 2.567 | 1.273 | 1.155 | 1.042 | 1.518 | 1.796 |
| **Y RNA** | 0.676 | 0.404 | 0.627 | 0.903 | 0.487 | 0.993 | 0.673 |
| **lncRNA** | 0.207 | 0.255 | 0.169 | 0.184 | 0.153 | 0.241 | 0.300 |
| **miRNA** | 0.177 | 0.251 | 0.146 | 0.172 | 0.135 | 0.224 | 0.207 |
| **rRNA** | 0.120 | 0.177 | 0.087 | 0.092 | 0.075 | 0.130 | 0.097 |
| **Vault RNA** | 0.001 | 0.000 | 0.001 | 0.001 | 0.001 | 0.001 | 0.002 |
| **snRNA** | 0.003 | 0.003 | 0.002 | 0.002 | 0.002 | 0.004 | 0.004 |
| **snoRNA** | 0.051 | 0.040 | 0.044 | 0.051 | 0.032 | 0.060 | 0.088 |

We analyzed the length of the reads aligned and assigned to each respective sncRNA category to determine if our annotation pipeline performed well and to describe the subspecies of various sncRNAs (**Table 1**). Notably, there was strong consistency in profiles among the six individual male rats (**Table 2**). The two canonical categories with known read lengths were checked first (miRNA and piRNA) and are discussed next.

### Few miRNAs are loaded into rat caput EPs compared to mice

The average read length for miRNA (22 nt) was exactly as expected [51,52] and showed a 90-percentile range of 19 – 27 nt (**Figure 3**). They account for only 0.18% of all reads. By contrast, miRNAs account for ∼60% of caput EVs in mice [34]. We set a cut-off of 10 reads per million (RPM) to determine which miRNA were loaded into caput EPs and identified 15 miRNAs, ten of which have previously been shown to exist either in sperm or EVs in various models, and five of which are uncharacterized miRNAs (**Table 3**). This is a far fewer number than those in mouse EVs (∼350) using a similar threshold [20]. Of the 15 miRNAs identified in rat, 7 overlapped with those reported in mice (miR-143, let-7c, let-7i, miR-26a, miR-99a, miR-143, miR-148 [20]). The most abundant miRNA we identified in rats, miR-184, did not meet the threshold for abundance in mice [20]. MiR-143, which is a hallmark of EVs in mice [20] and humans [34], was also found in rat caput EVs.

**Table 3.** Fifteen miRNA were identified as having substantial (> 10 RPM) expression in rat caput EVs, 10 of which are annotated and 5 that are putative. This is far fewer than expected based on mouse experiments [34]. The majority of miRNAs identified have previously been reported in either sperm or EVs, except for miR-1b. A miRNA considered a hallmark of caput EVs (miR-143) was also identified in our rat data set.

| Name | Reads per Million | Found in Sperm | Found in EVs |
| --- | --- | --- | --- |
| *miR-1b* | 16.2 | -- | -- |
| *miR-7a-1* | 19.7 | [106] | [107] |
| *miR-7a-2* | 19.9 | [106] | [107] |
| *miR-let-7c* | 10.0 | [106] | [107] |
| *miR-let-7i* | 11.7 | [106] | [107] |
| *miR-26a* | 11.7 | [108] | -- |
| *miR-99a* | 87.8 | [109] | -- |
| *miR-143* | 29.5 | [22] | [22] |
| *miR-148a* | 93.0 | [106] | [107] |
| *miR-184* | 493.7 | [106] | [107] |
| *AABR07005004.2* | 16.0 |  |  |
| *AABR07029907.2* | 34.7 |  |  |
| *AABR07057421.1* | 11.9 |  |  |
| *AABR07064716.2* | 34.2 |  |  |
| *AABR07064724.1* | 22.1 |  |  |

### piRNAs are abundant and coincide with 5’ tRNA fragments

The average read length for piRNA was also as expected (30 nt; [52]) with a 90-percentile range of 26 – 32 nt (**Figure 3**). We identified 81 piRNA loci that were substantially (> 10 RPM) expressed across the genome, with clusters on chromosome 1, 2, 10, 13, 14 and 17. piRNAs represented a substantial proportion (18.1%) of EPs in the rat caput epididymis. This is contrary to what is found in mouse models (< 0.05%; [34,35]) but not entirely unexpected as rat pachytene sperm are densely populated with piRNAs [53] depending on the stage of development [54,55] presumably to control the expression of transposable elements in the germline [56] by directing the catalyzation of *de novo* DNA methylation to suppress their expression [57]. The reasons for this discrepancy between rats and mice could be due to a number of reasons. First, this may represent a fundamental difference in the reproductive biology between the two species. Second, it may be because the piRNAs that do exist in mouse caput EVs are typically categorized as tRNA-halves instead of piRNA. We found that approximately 60% (49/81) of the expressed piRNA loci overlapped with known tRNA loci; these accounted for a significant portion of all piRNA annotated reads (82.3%). These are a known subclass of piRNAs termed tRNA-derived piRNA (td-piRNAs; [58]) so we further analyzed their origin and determined that virtually all of these reads aligned with the 5’ end of tRNA loci (98.98%). tRNA derived-piRNA (td-piRNA) have been described in the testes of marmosets where Piwi proteins were found to bind reads mapping to the 5’-tRNA^Glu^, 5’-tRNA^Gly^, and 5’-tRNA^Val^ loci [59]. Therefore, we categorized the tRNA type from which piRNAs were generated and determined that the majority were derived from either 5’-tRNA^Gly^ (53.3%) or 5’-tRNA^Glu^ (32.6%) loci; 5’-tRNA^Lys^ (8.5%) and 5’-tRNA^Val^ (2.3%) originating reads were also identified. It is intriguing that these three identified td-piRNAs account for three of the four td-piRNAs found here. To our knowledge this is the first evidence that td-piRNA are in a position poised to be delivered to sperm in the epididymis via EVs. Additionally, we append 5’-tRNA^Lys^, the third most abundant td-piRNA in our analysis, as a potentially significant td-piRNA in rats.

### tRNA fragments (tRFs) dominate the cargo of rat caput EPs

We then analyzed other sncRNA categories where read lengths can be variable depending on how the RNA is processed. In mouse models, nascent sperm in the caput contain few tRFs but their abundance gradually increases as sperm transit to the cauda [22,60], a pattern that is mirrored by the EVs found in the same epididymal compartment [34]. By comparison, reads from tRNA loci dominated the cargo in our rat caput EPs (∼79%). The reads aligned to tRNAs had an average read length of 30 nt and a 90-percentile range of 28 – 31 nt (**Table 1**), suggesting that they are either tRNA fragments (tRFs) or tRNA-halves, which are 14 – 30 nt and 30 – 40 nt in length, respectively [61].

We further categorized tRNA reads by aligning them to either the 5’ or 3’ end of a given tRNA locus and discovered that 99.35% of all reads aligned to the 5’ end (**Figure 4A**). There are two possible explanations for the overrepresentation of 5’-tRFs. First, rat caput EPs may be selectively loaded with 5’-tRF fragments; these have actions similar to RNA interference as governed by miRNAs, which act through the Argonaute pathway [62] to silence endogenous retroelements in embryonic stem cells and embryos [22]. Alternatively, our observation may be due to a technical bias in the sequencing library preparation procedure [61]. Libraries are amplified with PCR during preparation, which can be prematurely aborted if an RNA molecule contains modified tRNA nucleobases that would be too short for sequencing, thereby excluded, and not detected during analysis. In order to confirm the dominating presence of 5’-tRFs in rat caput EPs, a specific analysis pipeline (streamlined platform for observing tRNA – SPOt) would need to be used to prevent observation bias [63].

**Figure 4.**
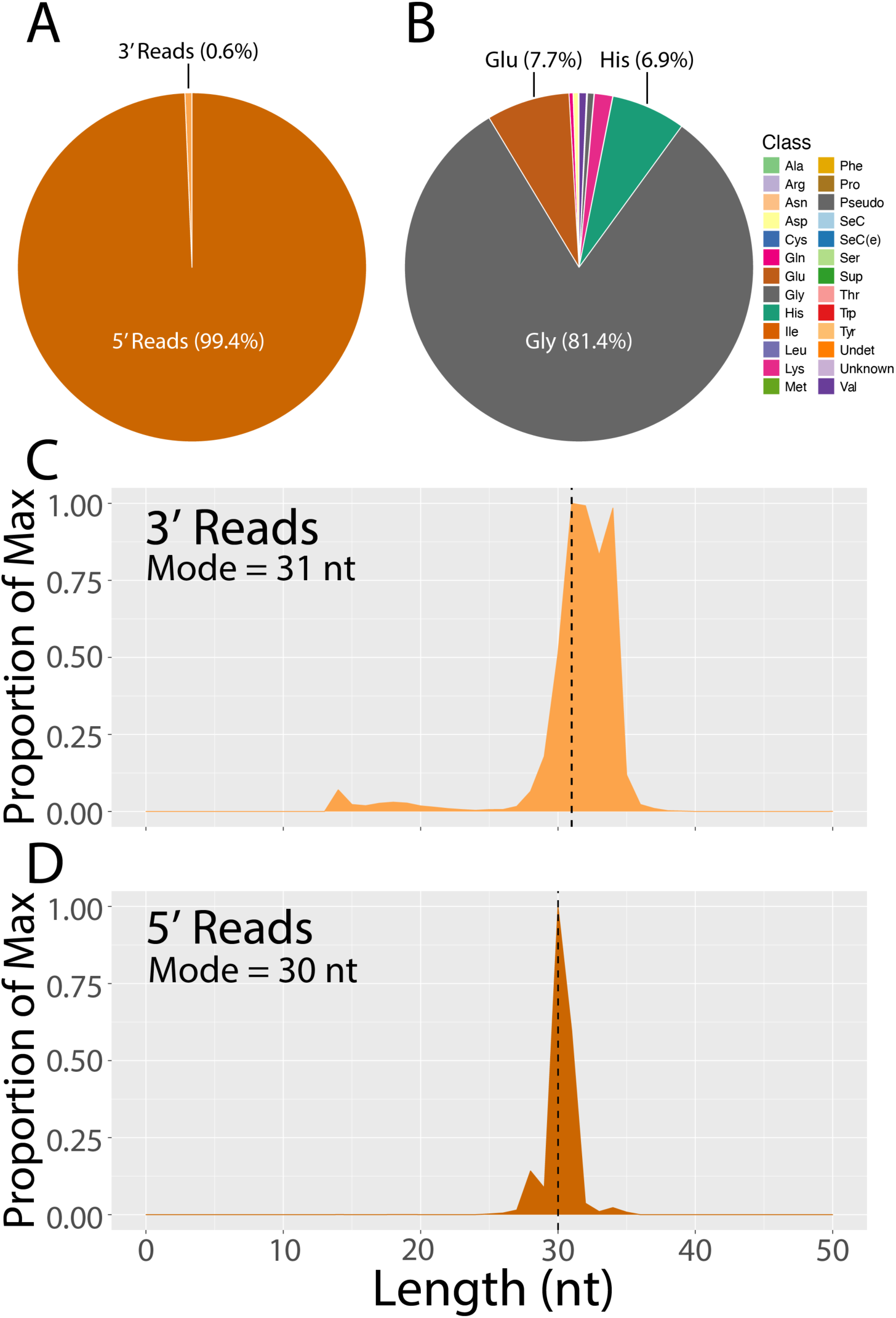
Detailed analysis of tRNA reads is shown. **A)** Reads align to the 5’ end of tRNA loci almost exclusively. **B)** 5’-tRNA^Gly^ is the primary tRNA fragment found in rat caput extracellular vesicles, followed by a substantially smaller proportion of 5’-tRNA^Glu^ and 5’-tRNA^His^. **C-D)** The size distribution of 3’ and 5’ tRNA reads are shown, respectively. Reads from the 3’ end of tRNA are slightly longer and have a broader distribution.

Given their size distribution, we were able to further categorize the 5’-tRFs reads, which show a 28 – 30 nt size distribution indictive of tRF-5c fragments, the longest of the three known 5’-tRFs, or tRNA-halves [64]. Alignment of reads to the tRNA structure (tRNAvis) confirmed their identity as tRF-5c fragments because reads did not align to the anti-codon stem loop, which would be characteristic of 5’ tRNA-halves [61]. Because of the significant overlap of reads identified between piRNAs and tRNA loci (presented above), we analyzed the proportion of tRNA reads that could be accounted for by piRNA loci. We found that the majority of tRNA reads (∼81%) were unique to tRNAs and could not be accounted for by overlapping piRNA loci.

Finally, we analyzed the amino-acid feature that tRNA reads were derived from and found that there was an over-representation of reads derived from Glycine-tRNAs (81.4%). Reads were also found to originate from glutamate (7.7%), histidine (6.9%), and lysine (1.7%) tRNA loci across the genome (**Figure 4B**). Finally, the size distribution for 3’ tRNA reads (**Figure 4C**) was similar to 5’ tRNA reads (**Figure 4D**).

### rRNA fragments are present in three distinct lengths

rRNA fragments (rRFs) have long been considered RNA degradation or apoptotic by-products and are generally disregarded for any functional value. Compared to miRNA, piRNA, and tRFs, little is known about the function or categories of rRFs and they have only recently been described [65] with nearly no information existing in rats that we could find. Interestingly, rRFs appear in immunoprecipitations with the Argonaute complex in mice and humans, which suggests a role for translational regulation [66]. In humans, three distinct categories of rRFs are known with differential expression in each of the rRNA categories (i.e., 5s, 5.8s, 18s, and 28s): short (18-21 nt), intermediate (24-25 nt), and long (26-33 nt) [65].

Here, we identify the presence of rRFs in rat caput EPs. Reads aligned to rRNA loci had an average length of 22 nt and a broad 90-percentile range of 17 – 31 nt (**Table 1**). rRNA is expressed as a 45s pre-rRNA, which contains the transcripts for 18s, 5.8s, and 28s rRNA, organized as cassettes of tandem repeats on the short arms of chromosomes 3, 11, and 12 of the rat genome [67]. The rat reference genomes are poorly annotated for these repeating cassettes because the number of repeats often varies between individuals. Hence, these regions are typically masked in alignment reference genomes. Nonetheless, there are a number of 5s rRNAs, which are expressed separately from the 45s pre-rRNA, and 5.8s rRNA loci that are annotated in the rat reference genome that we analyzed. Approximately 80% of rRNA derived reads aligned to 5.8s loci (**Figure 5A**). Subcategories of rRFs have been described from human samples where short (18-19 nt), intermediate (24-25 nt), and long (32-33 nt) rRFs are expressed depending on the rRNA type (e.g. 5s vs 5.8s) [65]. Our data generally fit the categories described in human samples: 5s rRNA resulted in either short or long transcripts (**Figure 5B**) whereas 5.8s rRNA resulted in intermediate fragments (**Figure 5C**). rRFs are sparsely reported in reproductive tissue (Humans, [68,69]; Bovine, [70]). We believe this is the first report of their presence in caput EP samples, meriting future studies on a role for rRFs in reproduction.

**Figure 5.**
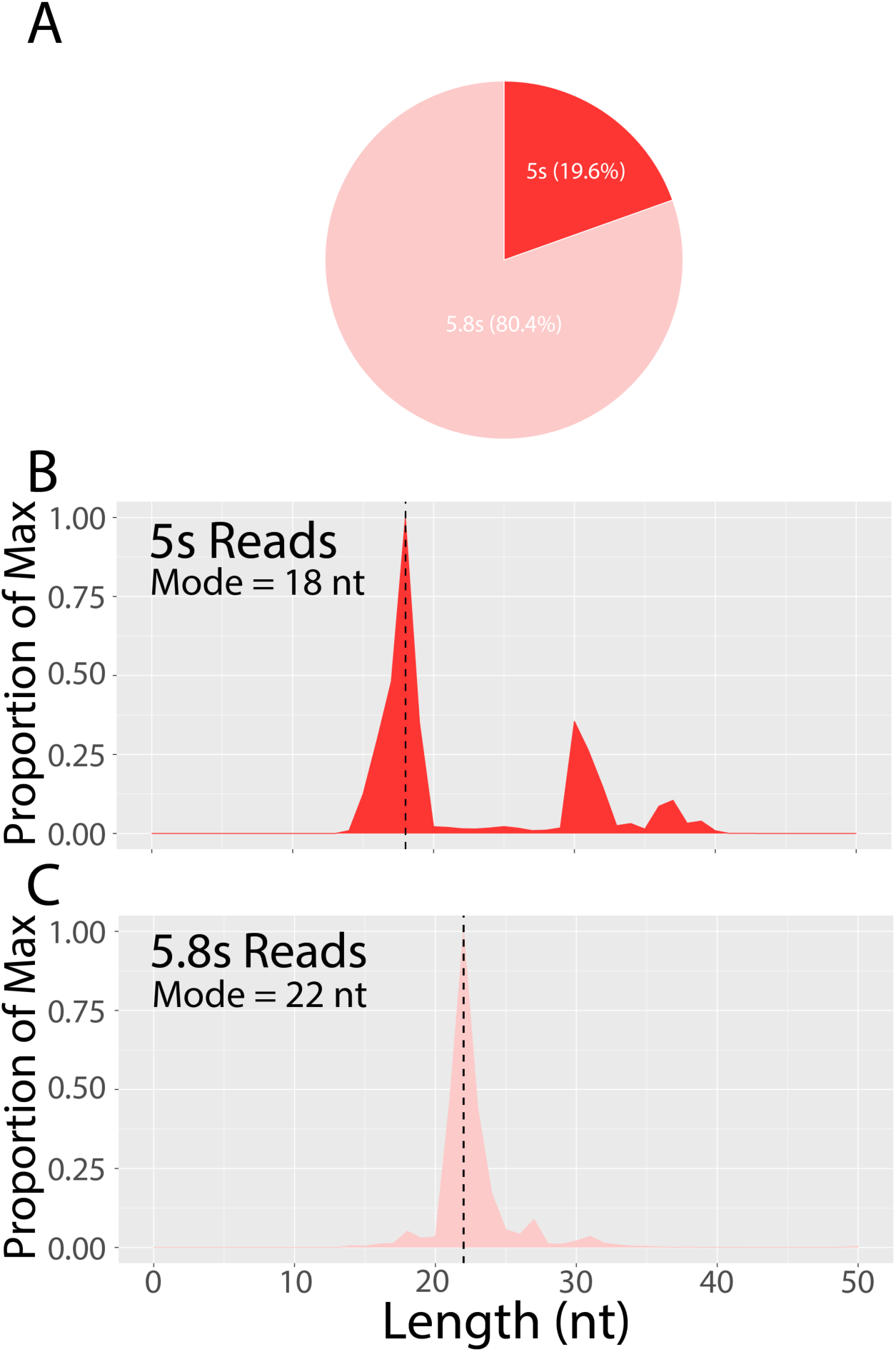
Reads that aligned to annotated rRNA loci are shown by subcategory. The rat genome is poorly annotated for rRNA, so the available 5s and 5.8s loci were used for categorization. **A)** The majority of rRNA reads aligned to 5.8s rRNA. **B-C)** The size distribution for reads aligned to 5s and 5.8s rRNA show profiles indicative of rRFs that are distinct from one another. **B)** 5s rRFs show a bimodal distribution with peaks at 18 and 30 nt. **C)** 5.8s rRFs show a single peak at 22 nt.

### Fragments of other RNA types are also present in rat caput EPs

Finally, we observed that the Y RNA was present in caput EPs at levels ∼4 times that of miRNA, which is surprising because we are not aware of any reports of Y RNA in epididymal EVs, although they are found in EVs of other tissue types [71–73]. Full length Y RNAs are ∼80 – 110 nt in length [74]. We identified 13 (of 28) Y RNA loci with substantial expression (> 10 reads per kilobase million – RPKM). These reads demonstrated a sharp peak at 31 nt and a smaller peak at 22 nt with a 90-percentile range of 22 – 31 nt (**Table 1**). These results suggest that rat caput EVs include fragments of Y RNA as a part of their sncRNA repertoire.

### Small RNA molecules are expressed from within CpG islands

We identified dense clusters of expression that appeared to be contained within the boundaries of GC-rich CpG islands (**Figure 6A**). CpG islands are typically annotated based on sequence characteristics defined as longer than 200-bp with a GC content higher than 50% and an expected-to-observed ratio greater than 0.6 [75]. Many of the first CpG islands were identified at the 5’ end of “housekeeping” genes [76,77] but have since been computationally and experimentally predicted across the genome, including in inter- and intragenic space [78]. Parts of CpG islands are sometimes transcribed on the 5’ or 3’ end of expressed genes and removed during splicing unless they extend into an exon, including within sncRNAs [78,79]. Because small RNAs have not previously been ascribed to the boundaries of CpG islands, we meticulously characterized these features, and further challenged this result by trying to ascribe reads contained with CpG islands to other surrounding or overlying genomic features.

**Figure 6.**
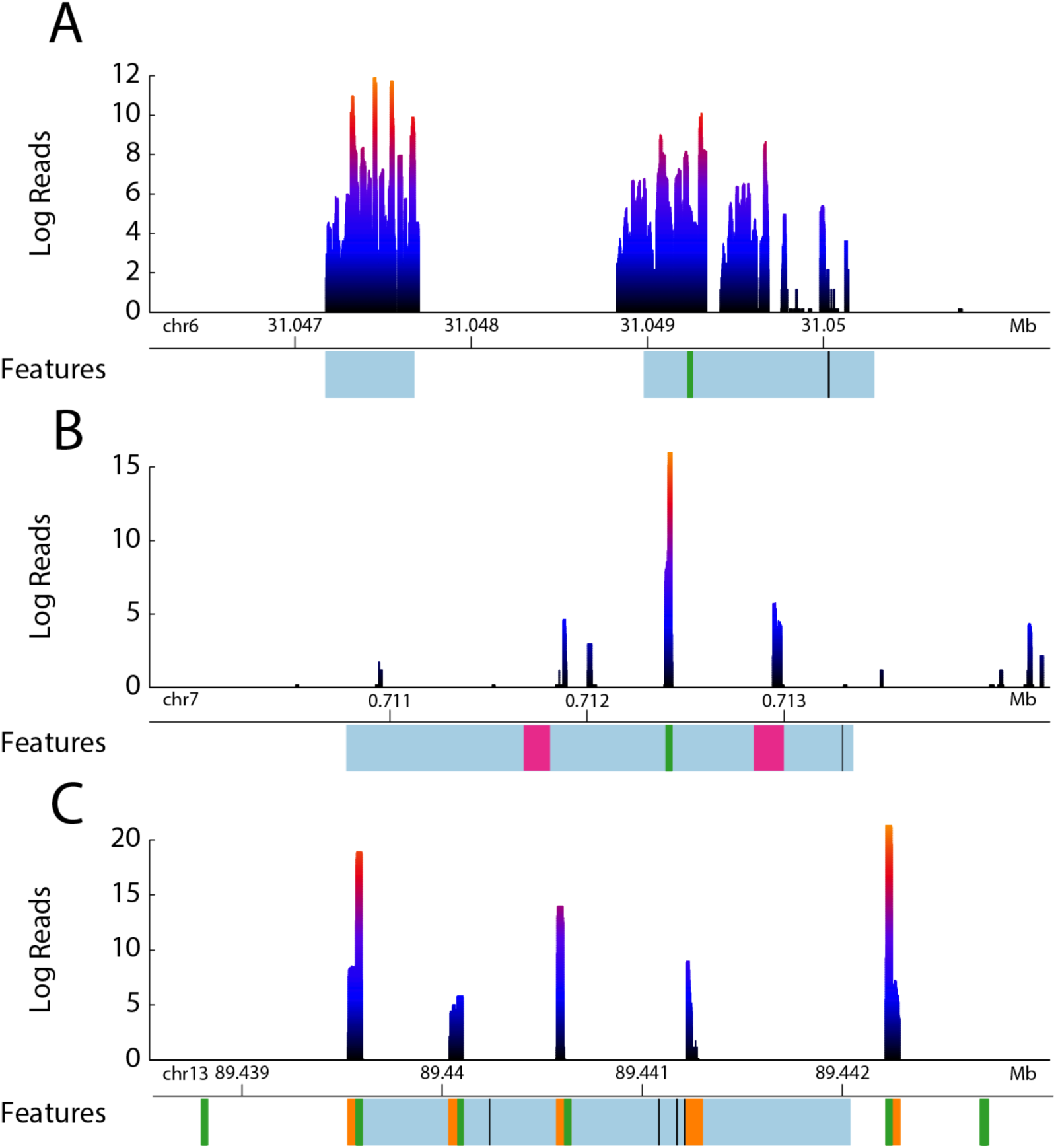
The log transformed density of reads is shown across three CpG islands with different characteristics. Feature annotations are shown below the x-axis and are indicated by color as follows: Light blue designates a CpG island; Green = piRNA, Black = predicted transcriptional start sites, Pink = 5.8s rRNA, and orange = tRNA. The density of reads is indicated by a heat map from black (minimum) to red (maximum) scaled to each respective y-axis. Plots were generated from BedGraph files with R (v4.2.0 [104]) and the Sushi package (v1.34.0, http://bioconductor.riken.jp/packages/3.1/bioc/html/Sushi.html [105]). **A)** Reads are shown expressed from within the boundaries of a CpG island on chromosome 6. The reads within the 3’ CpG island were not associated with the piRNA (green), while the 5’ CpG island had no other overlying features. **B)** Reads were identified within a CpG island on chromosome 7 but the majority of them aligned to a piRNA (green) and rRNA (pink) feature. **C)** Reads were identified within a CpG island on chromosome 13 but they aligned exclusively to piRNA and tRNA loci and are demonstrative of their typical expression profiles. The CpG islands from **B & C** (and all other CpG islands such as these) were removed from the analysis of small RNAs expressed from CpG islands. The two in **A** are representative of those that were used for further analysis.

In total we identified 50 CpG islands with significant expression (>10 RPKM) of small RNA molecules. We inspected the distribution of reads (SeqMonk) within each of these CpG islands and were able to rule out the majority of them due to overlapping features that had canonical expression patterns expected from features like rRNA and piRNA (**Figure 6B**) or tRNA (**Figure 5C**). These loci were removed from further analyses and the remaining 12 CpG islands were treated as an annotation category for sncRNAs. We found this category to be the third most abundant behind tRNA and piRNA reads (**Figure 3A** and **B**), and that the reads derived from CpG islands had a length distribution pattern that was distinct from all of the other analyzed categories (**Figure 3C**) with a median length of 19 nt and a 90-percentile range of 15 – 28 nt (**Table 1**). As expected, the GC content of reads aligned to CpG islands was substantial (∼78%) compared to the average of all aligned reads (∼50%).

We calculated a signal-to-noise (s/n) ratio to determine if the reads we observed within CpG islands were due to random chance [65]. When compared to the number of reads per kilobase million (RPKM) over a 10 kb rolling window across the entire genome, our identified CpG islands had a 609 s/n ratio. When compared to all CpG islands across the entire genome, our identified CpG islands had a 92 s/n ratio. Put simply, the reads contained within the identified CpG islands were 609 times more abundant than a random 10kb window in the genome and 92 times more abundant than an average CpG island, indicating that the reads we observed were very unlikely to be due to random chance.

Because CpG islands are not known units of small RNA expression, we then systematically considered whether the observed reads were due to other overlying features. We observed that those CpG islands that contained more piRNA loci appeared to be associated with more reads (**Figure 7A**). The function of piRNA is RNA-guided silencing of transposable elements, particularly in the germline before exit from the testis [54,80–82]. Hence, the presence of piRNA in caput EVs would not be surprising. However, because the read distribution we identified for other piRNA loci (e.g. **Figure 6B**) was very different from reads identified for CpG islands (e.g. **Figure 6A**) we considered that the transcripts we were observing might be secondary piRNAs generated via the ping-pong cycle wherein piRNAs are amplified by binding expressed mobile elements [54,83] or byproducts from their generation [84–86]. We therefore analyzed the relationship between piRNA loci and the reads expressed within CpG islands; when normalized for the read depth and length of the CpG island (RPKM) the relationship was virtually non-existent (R^2^ = 0.077, **Table 4**). We quantified the reads that aligned under known piRNA loci and found that ∼48% of the total reads within the boundaries of CpG islands were also associated with piRNA loci. However, we are hesitant to classify these reads as piRNA because they do not follow the canonical length distribution associated with primary or secondary piRNA (∼30nt) identified here and as reported elsewhere [87].

**Figure 7.**
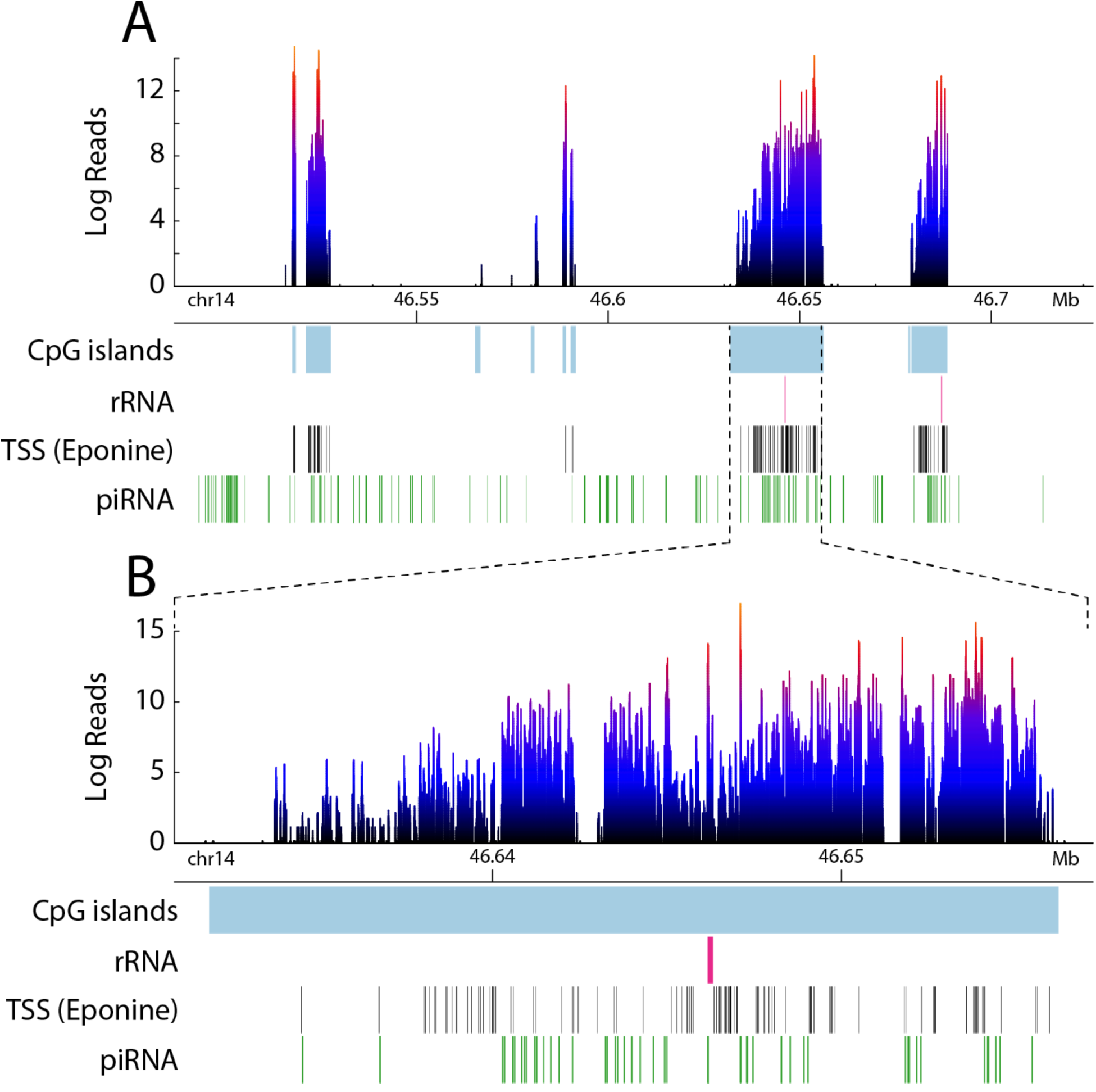
The log transformed reads from a cluster of 8 CpG islands on chromosome 14 are shown with the primary overlying features indicated by color and label below the x-axis. Light blue designates the CpG islands. Pink = 5.8s rRNA, Black = predicted transcriptional start sites, Green = piRNA. The density of reads is indicated by a heat map from black (minimum) to red (maximum) scaled to each respective y-axis. Plots were generated from BedGraph files with R (v4.2.0 [104]) and the Sushi package (v1.34.0, http://bioconductor.riken.jp/packages/3.1/bioc/html/Sushi.html [105]). **A)** The boundaries of the reads shown are clearly demarcated by CpG islands and could not be attributable to overlying features such as rRNA or piRNA. **B)** A detailed view of the largest CpG island in the chromosome 14 cluster. The reads are not associated with either the rRNA or piRNA features within the CpG island and are unique from those found elsewhere (Figure 5B & C).

**Table 4.** The genomic coordinates (Rnor v6 – Chromosome, Start Location, End Location) of 12 CpG islands found to have substantial expression (> 10 RPKM) of ∼19 nt small RNA transcripts are shown along with the length of the CpG island, raw read count, and length-corrected read count (RPKM) found within each. The number of features found overlapping each CpG island is also shown.

| Chromosome | Start Location | End Location | Length (nt) | Read Count | RPKM | piRNA Loci | pRNA Protein Loci | rRNA Binding Protein Loci | rRNA Loci |
| --- | --- | --- | --- | --- | --- | --- | --- | --- | --- |
| Chr 1 | 11,903,555 | 11,916,838 | 13,283 | 112,070 | 63.16 | 19 | 1 | 0 | 1 |
| Chr 1 | 11,961,297 | 11,975,108 | 13,811 | 278,633 | 151.02 | 27 | 1 | 0 | 1 |
| Chr 5 | 91,124,274 | 91,141,016 | 16,742 | 341,585 | 152.72 | 26 | 1 | 0 | 1 |
| Chr 6 | 30,627,068 | 30,633,859 | 6,791 | 177,949 | 196.15 | 13 | 0 | 0 | 2 |
| Chr 6 | 31,047,175 | 31,047,676 | 501 | 10,347 | 154.59 | 0 | 0 | 0 | 0 |
| Chr 6 | 31,048,983 | 31,050,286 | 1,303 | 3,282 | 18.85 | 1 | 0 | 0 | 0 |
| Chr 14 | 46,517,594 | 46,518,401 | 807 | 113,831 | 1,055.86 | 1 | 0 | 0 | 0 |
| Chr 14 | 46,521,126 | 46,527,599 | 6,473 | 170,612 | 197.30 | 10 | 0 | 1 | 0 |
| Chr 14 | 46,588,206 | 46,589,034 | 828 | 39,955 | 361.21 | 0 | 0 | 1 | 0 |
| Chr 14 | 46,590,294 | 46,591,521 | 1,227 | 2,562 | 15.63 | 1 | 0 | 1 | 0 |
| Chr 14 | 46,631,880 | 46,656,213 | 24,333 | 429,235 | 132.04 | 44 | 0 | 1 | 1 |
| Chr 14 | 46,679,234 | 46,688,547 | 9,313 | 160,403 | 128.93 | 17 | 0 | 0 | 1 |

The byproducts of secondary piRNA generation are typically ∼19-nt, which matches the length distribution we see here. We consider this the most likely possibility of those considered, but our observation does not fully match their description. First, a subset of the CpG islands that we identified were not associated with any known piRNA annotations. Second, the mechanism by which piRNA biproducts are generated usually results in 19-nt RNA from both strands and broad areas that are only constrained by the targets that primary and secondary piRNA bind. We therefore analyzed the source strand of the identified reads and found they are almost exclusively expressed from a single strand in the orientation of the primary piRNA annotations in the instances there were any at all (**Table 4**). Third, we looked at overlying repeat elements (LINE, SINE, and LTRs) and found these CpG islands void of any such annotated repeats. Because piRNA and tRNA are often associated, we also analyzed the proportion of reads aligning to tRNA loci and found none of the reads (0%) within CpG islands were expressed from known tRNA loci. In a subset of the CpG islands we observed, there was an abundance of predicted (eponine) transcriptional start sites (**Figure 7B**), but this was not always the case (e.g. **Figure 6A**).

Finally, half (6/12) of the identified CpG islands were associated with annotated 5.8s rRNA loci (**Table 4**). None of the CpG islands we identified were in proximity to known locations of rRNA cassettes, namely, on the short arms of chromosomes 3, 11, and 12 [67]. Notwithstanding, the rat genome is poorly annotated for rRNA, so to overcome this shortcoming in the available annotations we used BLAST to align the raw sequences expressed from CpG islands to the available rat rRNA sequences: 45s, 32s (which includes the 28s rRNA and 5’ sequence between the transcription start site and 28s), 28s, 18s, and 5.8s. Approximately one third (∼34.3%) of all reads derived from CpG islands aligned to some form of rRNA or precursor rRNA. Of those reads that aligned to any form of rRNA, the vast majority were derived from the 28s sequence (86.1%), while the 18s (11.15%), and 5.8s (0.16%) accounted for a small proportion of the reads. The remaining reads (2.6%) aligned to the external or internal transcribed spacers within the full length 45s rRNA precursor transcript. Given the proximity of these reads to apparent rRNA cassettes, we cannot rule out that some portion of the observed CpG islands may be a part of the ∼30 kb non-transcribed spacer that separates 45s repeats. But, as the name implies, these portions are up or downstream of rRNA and should not be transcribed.

In summary, while it appears that the CpG islands we identified as expressing 19 nt small RNAs were in the vicinity of heretofore unannotated rRNA or piRNA loci, the reads observed expressed within CpG islands cannot be fully accounted for by either of these designations nor are their expression profiles congruent. Therefore, we believe this is a novel category of small RNA that warrants additional investigation.

## Discussion

Epididymosomes from the caput are an essential component of sperm maturation and acquisition of function [5,88,89]. The proteins transferred to sperm via caput EVs are believed to assist in sperm-zona binding [16] and the development of motility in sperm [90]. Caput EVs are also implicated in the control of heritable non-genetic phenotypes, as their sncRNA cargo is altered by stress [22,26,29], diet [27,30,91], and alcohol consumption [31]. Thus, caput EVs and their contents play roles in development and the health and behavior of offspring. Support for the role of sncRNA in transgenerational inheritance is bolstered by experiments in which they are directly injected into zygotes or sperm [32]. This method has been used to demonstrate inter- or transgenerational inheritance of metabolic [27,30,92] and behavioral phenotypes [93–95].

Here, we provide the first comprehensive characterization of small RNAs derived from caput EPs in the rat, and demonstrate quantitative and qualitative aspects unique to this species. EPs isolated from the caput epididymis match the size range of epididymosomes and their small RNA contents are dominated by tRFs and piRNA, containing far fewer miRNAs than expected from other organisms (Mice, [29,34,35]; Humans, [96]). We also identify Y RNA fragments in caput EPs for the first time in any organism, although they are expressed in EVs of other organs [71–73]. Finally, we identify a potentially novel small RNA molecule that is expressed from GC-rich CpG islands that cannot be accounted for by known overlying small RNA features, and which have a unique size distribution that is distinct from other small RNAs analyzed.

The small RNA contents of EVs from the mouse caput epididymis are primarily miRNAs (∼60% - [29,34,35]), with over 350 expressed [20]. This is in contrast to the rat in which miRNAs were a small portion (0.18%) of the small RNA complement. There were also substantial species differences in tRFs, which are the second most abundant category (∼30%) in mouse caput EVs while piRNAs are present at very low levels in mice (<0.05% - [34,35]). Here, we show in rats that tRFs are by far the most abundant category (∼79%) and piRNAs are second most abundant (∼18%). Further work is needed to validate these findings and to understand why there are these dramatic species differences in the EV sncRNA cargo.

Finally, we believe we have discovered a novel type of small RNA that is expressed from within the boundaries of CpG islands. Based on the analyses we performed and the data presented here, we do not believe that alternative sncRNAs (rRNA fragments, piRNAs, or piRNA amplification byproducts) can account for these 19-nt small RNAs. More specifically, these “CpG island small RNAs” extend outside of the boundaries of annotated rRNA and piRNA, which account for only a portion of the observations, and do not abide by their respective canonical length characteristics. It is possible that these observations could be the 19mer byproduct of piRNA amplification observed elsewhere [84–86] but there are 3 reasons we do not think this is likely. First, we observe these reads in EPs while the byproducts of piRNA amplification have only been found to exist in pre-pachytene spermatocytes and should be derived from both the sense and anti-sense strands, a characteristic we do not see. If this observation is due to 19mer piRNA amplification byproducts generated in the epididymis instead of spermatocytes themselves, our data would suggest that secondary piRNA are produced in the epididymis and they, along with their byproducts, are actively transported to spermatozoa transiting the epididymis. Second, the reads we observe are expressly within the boundaries of CpG islands with a GC content not reported elsewhere. The specificity of the expression loci, and the lack of a high GC content in other reports on the production of secondary piRNA make their identity as byproducts unlikely. Finally, we observe expression from multiple CpG islands not linked with known piRNA loci. If these are indeed the byproduct of piRNA amplification, our data would represent the identification of novel piRNA loci that would seem to be important for the final steps of sperm maturation. Mature sperm transiting the epididymis should be transcriptionally quiescent [97–100], and therefore transposons, which are the presumed target of piRNA, should not be expressed. Ascertaining a functional role of these small RNAs requires further investigation.

In summary, EPs from the rat caput epididymis carry a complex repertoire of small RNAs and their contents are substantially different from the mouse. The dominant features we identify are tRNA fragments and piRNAs derived from tRNA loci. MicroRNAs are poorly represented in stark contrast to mice. We also report the presence of two known types of small RNA not previously seen in caput EVs, rRNA fragments and Y RNA fragments, and identify a potentially novel small RNA we have termed CpG island (CpGi) sRNAs. Considering that epididymal EVs can transfer their RNA cargo to sperm, these data represent an exciting collection of possibilities for future research on mechanisms of basic reproductive biology and the complexities of epigenetic transgenerational inheritance.

## Acknowledgement

The authors recognize Mandee Bell and Lindsay M. Thompson for their assistance with animal husbandry, treatment, and sample collection. We thank Michelle Mikesh for her assistance in the use of transmission electron microscopy, and Dr. Axel Brilot and Dr. Evan Schwartz for their assistance in the use of cryo-electron microscopy. We also acknowledge the excellent work (small RNA library preparation and sequencing) performed by the Genomic Sequencing and Analysis Facility at UT Austin, Center for Biomedical Research Support. RRID#: SCR_021713. Finally, we would like to thank Dr. Nicholas Peppas and his laboratory for training, guidance, and execution of particle tracking analysis and Dr. Blerta Xhemalce for reviewing our approach and data.

## Grant funding

Supported by R35 ES035024 and R21 ES034067 to A.C.G., and PhRMA Foundation Postdoctoral Fellowship to R.G.

## Conflict of Interest

The authors declare no conflict of interest.

## Data Availability

The raw sequence data reported in this paper have been deposited in the Genome Sequence Archive [101] at the National Genomics Data Center [102] that is publicly accessible at https://ngdc.cncb.ac.cn/gsa accession GSA: CRA008039.

## ARRIVE

The study was carried out according to the ARRIVE guidelines [41].

## Notes

### Competing Interest Statement

The authors have declared no competing interest.

### Summary of Updates

Added new data shown in Figure 2 for electron microscopy and cryo-EM of extracellular particles. This work was done by Dana L Sheinhaus who was now added as a co-author.

